# Body site-specific and disease-specific virulome in the human microbiome

**DOI:** 10.1101/2020.12.13.403006

**Authors:** Fei Liu, Wanting Dong, Yaqiong Guo, Qian Xiong, Na Lu, Xiaofeng Song, Yong Xue, Demin Cao, Xinyue Fan, Yuan Fang, Zhiyuan Li, Jian Cao, Yanan Wang, Guowei Yang, George F. Gao, Fangqing Zhao, Baoli Zhu

**Author notes:** These authors contributed equally to this work (Fei Liu, Wanting Dong). Corresponding author: Baoli Zhu.

## Abstract

Human body habitats are home to a diverse array of microbes, and within these microbial ecosystems, there are exchanges of genetic material, including virulence factors (VFs). Little is known about the diversity and abundance of VFs in different body sites and different types of diseases. We developed a virulome analysis pipeline using the species-specific sequence identity inferred from intraspecies ANI values to precisely assign reads to virulence factors. We characterized the human virulome from four body habitats, including the gut, oral cavity, skin, and vagina. Specifically, the diversity and abundance of VFs in the oral cavity were significantly higher than those in other body sites, including stool. We highlight the importance of sex-specific analysis when studying the human virulome. We analyzed data from more than 4,000 samples across healthy and diseased subjects and 13 types of diseases from different metagenomic sequencing studies to characterize the disease-specific virulome. Atherosclerotic cardiovascular disease (ACVD) has a more diverse virulome than other diseases tested. Notably, many VFs, including genes for secretion systems and toxins, are more abundant in diseased subjects than in healthy controls. We present, to our knowledge, the most comprehensive healthy and diseased virulome dataset yet created.

## Background

The human microbiome has been identified as an essential factor in many diseases, including obesity^1^, type 2 diabetes^2^, and cirrhosis^3^. Microbial metabolites and components influence the susceptibility of the host to many immune-mediated diseases and disorders^4^. Pathogen colonization is controlled by bacterial virulence and through competition with commensals^5^. Virulence factors (VFs) are typically defined as pathogen components whose loss specifically impairs virulence but not viability, including adhesins, toxins, exoenzymes, and secretion systems^6^. They are produced by pathogens that could cause diseases^7^. Although nonenterotoxigenic *B. fragilis* (NTBF) is a common component of the colon, enterotoxigenic *Bacteroides fragilis* (ETBF), which secretes *B. fragilis* toxin, could induce colonic tumors^8^. Recent studies suggest that colorectal cancer (CRC) is influenced by *pks+ Escherichia coli*, which contains the colibactin-producing *pks* pathogenicity island, directly impacting oncogenic mutations^9,10^. These results highlight the need to characterize the microbiome at the strain level and the differences in VFs between healthy and diseased individuals. Moreover, we should also pay more attention to microbial communities for evaluating pathogenicity^11^. With metagenome sequencing, we can observe all microbial genes present in a complex community^12^, including VF genes. However, the extent and diagnostic implications of virulome contributions to different types of the disease remain unknown.

Currently, the virulence factor database (VFDB, http://www.mgc.ac.cn/VFs/) provides up-to-date knowledge of VFs from various bacterial pathogens. It serves as a comprehensive warehouse of bacterial pathogenesis knowledge, including a core dataset covering experimentally verified VFs^13^. There are also many other virulence factor databases, including Victors^14^, PATRIC^15^, and PHI-base^16^. Hidden Markov models^17^, deep convolutional neural network models^18^, and VFanalyzer^19^ are used for VF classification in bacterial genomes. Whole-genome sequencing is an effective method to comprehensively identify VFs. However, the reliable and efficient characterization of VFs in the metagenome remains a challenge. Biosynthetic gene clusters could be predicted using ClusterFinder^20^, which also yields false-positive results. We wish to apply a reasonable and stringent cutoff to the VF analysis to exclude potential false positive matches.

Here, we used species-specific sequence identity (SSI) inferred from the mean ANI values per species to precisely assign reads to virulence factors. As little is known about the abundances and diversity of VF profiles in different body habitats, we randomly selected 1,497 metagenome datasets from habitats within the human skin, oral cavity, gut, and vaginal from the Human Microbiome Project (HMP) cohort to carry out virulome analysis. We highlight the importance of sex-specific analysis when studying the human virulome. We analyzed data from 4,000 samples across healthy and diseased subjects and 13 types of diseases from different metagenomic sequencing studies to characterize the disease-specific virulome. We present, to our knowledge, the most comprehensive healthy and diseased virulome dataset yet created.

## Results

### Curation of the virulence factor database and establishment of the methodology for virulome classification

We curated the gene annotation of experimentally verified VFs in the VFDB, which comprises 3,228 experimentally verified gene sequences from 53 species of bacterial pathogens. *Legionella pneumophila, Escherichia coli*, and *Pseudomonas aeruginosa* were the top three species based on the number of their VF gene sequences in the dataset (Table S1). We manually inspected the VF gene categories. Adherence, T4SS, T3SS, invasion, toxin, and iron uptake systems were the top six categories (Table S2).

VFs are often species-specific and variably conserved between species^21^. The average nucleotide identity (ANI) was developed for bacterial species classification^22^. We performed intraspecies ANI analysis for each of the 53 species. Figure 1A shows that the ANI values range from 85.3% *(Pseudomonas stutzeri)* to 99.9% *(Bordetella pertussis)* for different species. We performed BLAST searches against the chromosome sequences in the complete bacterial genomes using species-specific sequence identity (SSI) thresholds and different fixed nucleotide identity cutoffs ranging from 99% to 90%. Barplot shows the number of pathogenic and nonpathogenic strains that hit at least one VF under different cutoffs (Figure 1B). In this experiment, SSI achieved almost the same high precision as 100% and 99% but at a markedly higher recall (Figure 1C). SSI performed the best in accuracy and F1 scores since it identified a high number of TPs and did not introduce many FPs.

**Figure 1.**
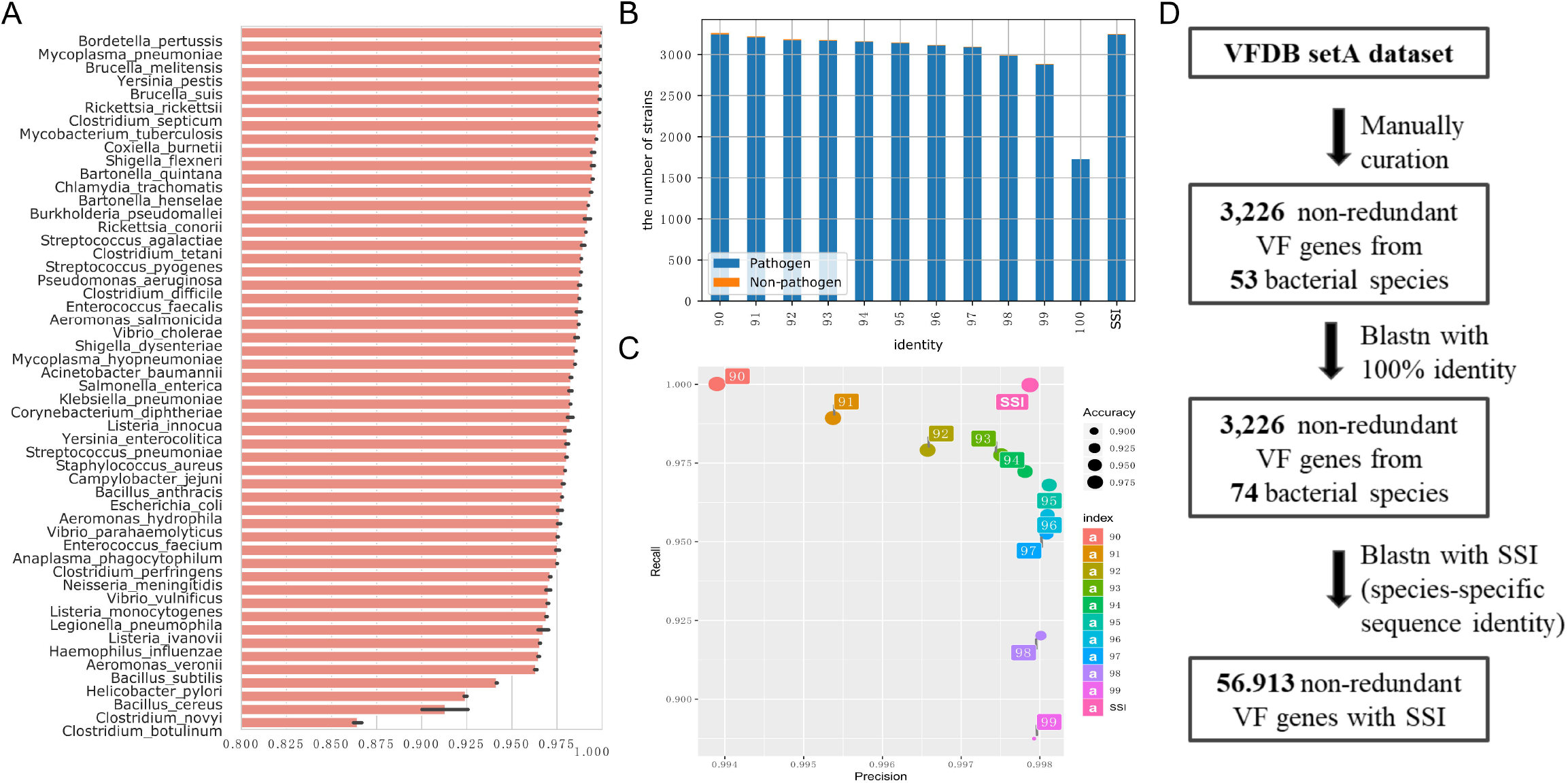
Comparison of the intraspecies whole-genome average nucleotide identity and accuracy of different thresholds for VF identification. (A) Barplot depicting the average nucleotide identity values of the 53 species of bacterial pathogens. (B) Barplot showing the number of pathogenic and nonpathogenic strains that hit at least one VF under different cutoffs. (C) Precision and recall graph for pathogenic and nonpathogenic strain identification under different cutoffs. We performed intraspecies ANI analysis for each of the 53 species. Figure 1A shows that the ANI values range from 85.3% *(Pseudomonas stutzeri)* to 99.9% *(Bordetella pertussis)* for different species. We performed BLAST searches against the chromosome sequences in the complete bacterial genomes using species-specific sequence identity (SSI) thresholds and different nucleotide identity cutoffs ranging from 99% to 90%. In this experiment, SSI achieved almost the same high precision as 100% and 99% but at a markedly higher recall (Figure 1C). SSI performed the best in accuracy and F1 scores since it identified high TPs and did not introduce many FPs. (D) Schematic representation of the curation of the VF dataset.

To further confirm our method’s accuracy, we compared the sequence identity of experimentally verified VFs between strains within one species to the mean ANI value in the species. Two experimentally verified VFs, namely, VFG005177 (gb|NP_664456) and VFG000959 (gb|NP_269190), were found in two strains, that is, *Streptococcus pyogenes* MGAS315 and *Streptococcus pyogenes* M1 GAS. The two genes’ sequence identity was 98.9%, which is very similar to the mean ANI (98.8%) of *Streptococcus pyogenes.* In addition, VF identification that relies on fixed criteria by loose cutoffs may result in misannotations. For instance, when using an 80% identity cutoff, the experimentally verified gene *east1* in *Escherichia coli* ONT:HND str. A16 can be found in many nonpathogenic strains, including the genome of *Candidatus Sodalis pierantonius* str. SOPE (CP006568.1). However, no experimentally verified virulence factor has been reported in this strain.

We identified a total of 2,893 VF gene sequences distributed across 5,250 strains within 74 species using a nucleotide identity cutoff value of 100% for the BLAST search against the chromosome sequences in the complete bacterial genomes. We manually inspected the newly identified species and found that all of them were also pathogens that could cause diseases, such as *Mycobacterium africanum*, *Klebsiella aerogenes*, and *Pseudomonas fluorescens*. This indicated that experimentally verified VFs were incomplete in the VFDB. In addition, we identified 31 prophage-encoded VFs, most of which were exotoxins.

We developed a virulome analysis pipeline that uses SSI inferred from the mean ANI values per species to precisely assign reads to virulence factors (Figure S1). With our expanded VF database termed VFGSSI, reference sequences of VFs were carefully chosen as seeds and integrated into the virulome analysis pipeline, making our database more comprehensive (Figure 1D). A list of pathogens in VFGSSI that can cause infections of the gastrointestinal tract or not and diseases they may cause are shown in Table S3 and Table S4.

### Different body sites have distinct virulomes

We analyzed 1,497 metagenome datasets from habitats within the human skin, oral cavity, gut, and vagina from the HMP cohort (Figure 2A). The overall alpha and beta diversity values for each body site were similar at the microbiome and virulome levels. The Shannon diversity values of the microbiome (Figure S2A) and virulome (Figure 2B) in the oral cavity were significantly higher than those in other body sites. Principal coordinate analysis of Bray-Curtis dissimilarities showed that the primary patterns of variation in the microbiome (Figure S2B) and virulome (Figure 2C) followed the major body sites (oral cavity, gut, skin, and vagina).

**Figure 2.**
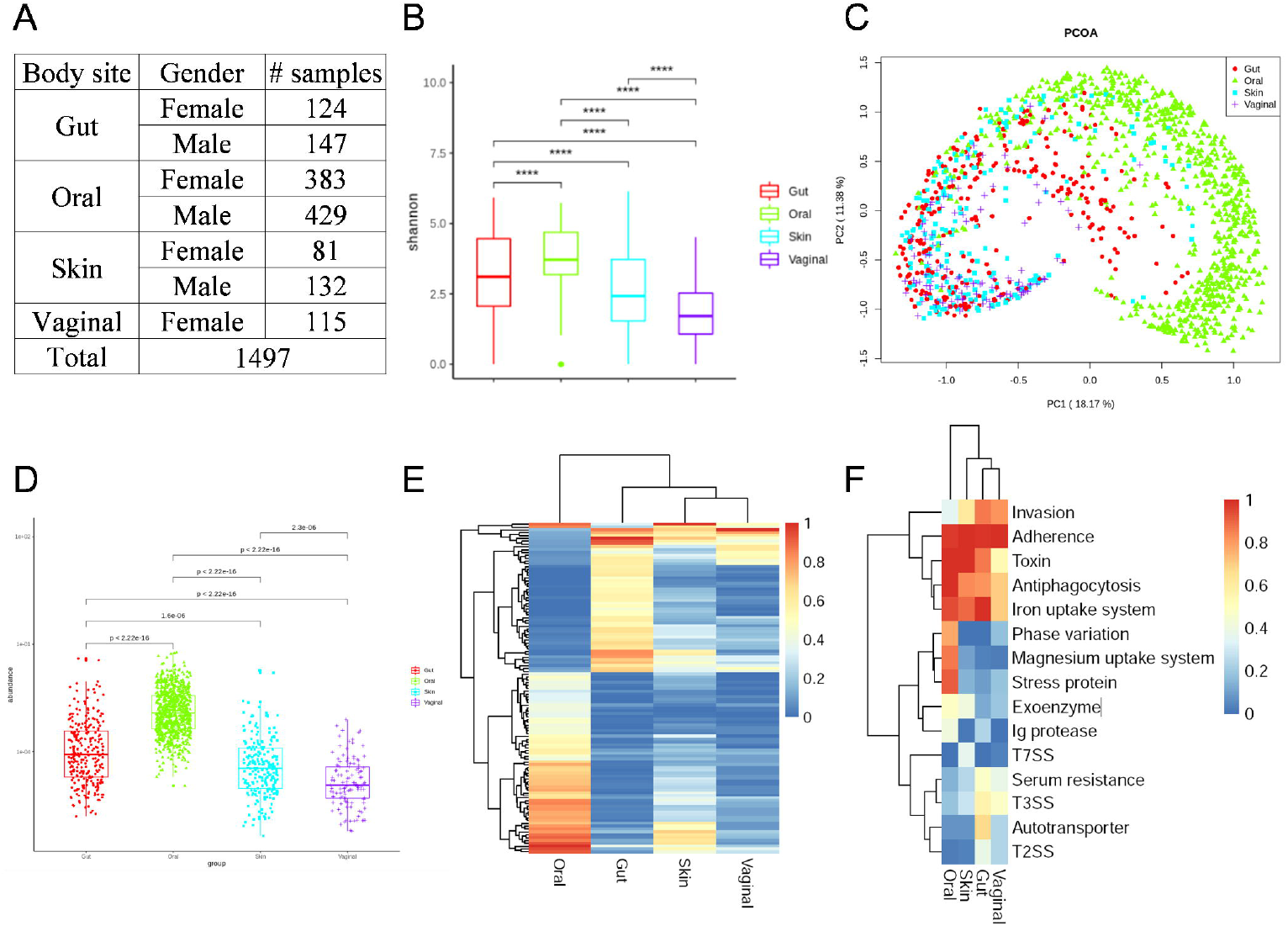
Different body sites have a distinct virulome. (A) Number of samples analyzed in the study. (B) Boxplot of the Shannon diversity indexes of all samples from different body sites based on VF abundance profiles. *p < 0.05, **p < 0.01, ***p < 0.001, ***p < 0.0001, Wilcoxon rank-sum test. (C) Principal coordinate analysis of Bray-Curtis dissimilarities showing the virulome. The first principal coordinate is shown by the x-axis, and the second principal coordinate is shown by the y-axis. (D) Comparison of the mean VF abundance. The centerline represents the median for each boxplot, and the boxes correspond to the 25th and 75th percentiles; all data points are shown. Hierarchical clustering of the prevalence of 106 VF genes (E) and 15 VF functional categories (F) that were hit in one of the body sites and are present in 20% or more of the samples in at least one body site. For the virulome analysis, the mean VF abundances in oral samples were significantly higher than those in other body sites. As expected, the vagina had the lowest total VF abundance. Additionally, the Shannon diversity values of VFs in the oral cavity and gut were significantly higher than those of VFs in other body sites.

A unique body site virulome composition was apparent. The mean VF abundances in the oral cavity were significantly higher than those in other body sites (Figure 2D). As expected, vaginal sites had the lowest VF abundance. Furthermore, the mean VF abundances in the samples at six major body sites are shown in Figure S3.

Specifically, the VF abundance in buccal mucosa was significantly higher than the VF abundance of other body sites. Hierarchical clustering of the prevalence of 106 VF genes (Figure 2E) and 15 VF functional categories (Figure 2F) is shown. In addition, we also performed LEfSe analysis to compare VFs (Figure S4). Specifically, in the oral cavity, the most differentially abundant VFs were capsular polysaccharide genes from antiphagocytosis.

The shared and unique VF genes among the groups were investigated. We found that 200 VFs were shared among body sites, accounting for 33.8%, 23.8%, 23.4%, and 43.8% of the total VFs identified in the gut, oral cavity, skin, and vagina, respectively (Figure S5A). Interestingly, the oral cavity and skin shared more VFs (689 types) than those shared between the gut and oral cavity (443 types) or between the gut and skin (444 types) (Figure S5B).

Interestingly, women showed a higher VF abundance in the skin and gut than men (ANOVA, p < 0.05, Figures S6A and S6B). Specifically, females had higher VF abundances in the anterior nares. In addition, sex-specific VFs for each body site were analyzed using LEfSe (Figures S7, S8, and S9). The availability of longitudinal samples of different body sites over two years from individuals who did not take antimicrobial drugs afforded us the ability to investigate the stability of virulomes over time (Figures S6C and S6D). There was no significant difference among samples from the same individuals except for the vagina, verifying that virulomes remained stable over a long period in different body habitats.

### Different disease types have distinct virulomes

We focused on 13 types of diseases for which the virulome is largely unknown, including colorectal carcinoma (CRC), atherosclerotic cardiovascular disease (ACVD), inflammatory bowel disease (IBD), obesity, hypertension, Parkinson’s disease (PD), non-small cell lung cancer (NSCLC), hepatocellular carcinoma (HCC), gastric cancer (GC), liver cirrhosis (LC), melanoma, renal cell carcinoma (RCC), and *Mycoplasma pneumoniae* pneumonia (MPP) (Figure 3A). As the original sequencing data of healthy individuals were missing in the NSCLC, RCC, melanoma, and HCC datasets, we developed an independent healthy cohort that served as a negative reference using the HMP gut data as mentioned above, which made intergroup comparisons possible.

**Figure 3.**
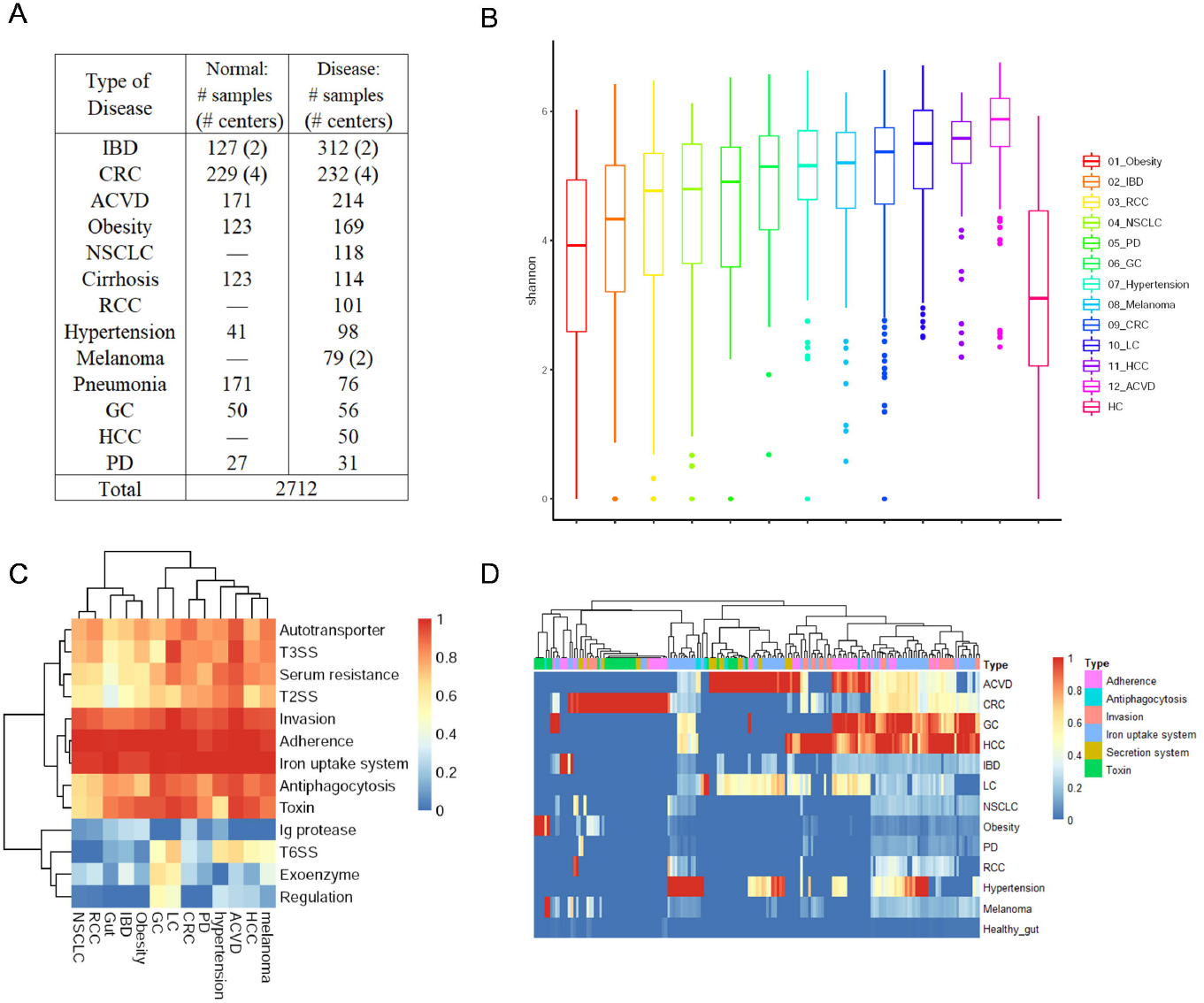
Different disease types have a distinct virulome. (A) Number of samples analyzed in the study. Dashes indicate data not available. ACVD, atherosclerotic cardiovascular disease; IBD, inflammatory bowel disease; CRC, colorectal carcinoma; NSCLC, non-small cell lung cancer; HCC, hepatocellular carcinoma; GC, gastric cancer; PD, Parkinson’s disease; RCC, renal cell carcinoma. (B) Boxplot of the Shannon diversity indexes of all samples from different types of diseases based on VF abundance profiles. (C) Hierarchical clustering of the prevalence of VF categories that were hits in one of the disease types and were present in 20% or more of the samples in at least one of the disease types. (D) Hierarchical clustering of the mean abundance of representative VFs for each type of disease. The top 10% (referring to the ratio of VF type numbers) of the most abundant VF types in each type of disease were considered the representative VFs.

First, we found that ACVD had a more diverse virulome than all the other disease types tested (P-value < 0.01 for each disease, Wilcoxon rank-sum test; Figure 3B). Compared to their own healthy controls, ACVD, CRC, and LC showed a higher diversity of VFs (p < 0.01, Figures S12, S13, and S14). In contrast, we did not find a more diverse virulome in obesity, IBD, PD, GC, and hypertension compared with their healthy controls.

Next, VF category prevalence was compared between diseases, and a disease-specific virulome composition was also clear (Figure 3C). We initially defined three groups for further VF category classification: high prevalence (>90%), medium prevalence (with prevalence ranging from 70% to 90%), and modest prevalence (<70%). VF categories including invasion, adherence, and iron uptake system composed the high prevalence group, characterized by consistently high prevalence in healthy and disease groups. Another six VF categories, including toxin, antiphagocytosis, autotransporter, T2SS, serum resistance, and T3SS, were the medium group members and were predominant in specific diseases. VF categories such as T6SS, Ig protease, exoenzyme, and regulation were divided into the modest group for their less predominant prevalence.

Moreover, hierarchical clustering of the mean abundance of representative VFs for each disease type is shown in Figure 3D. The top 10% (referring to the ratio of VF type numbers) of the most abundant VF genes in each type of disease, which were considered the representative VFs, are summarized in Supplementary Table S5. Specifically, compared to HMP healthy individuals, many VFs belonging to toxins were more abundant in obese individuals, while VFs encoding the iron uptake system were more abundant in hypertensive individuals. T6SS and antiphagocytosis genes were more abundant in patients with ACVD than in their healthy controls (Figure S15). Apart from invasion, adherence, and the iron uptake system, which were the universally discovered representative VF categories in those diseases, two clusters of VFs encoding secretion systems and toxins were found in ACVD and CRC patients, respectively, the existence of which distinguished CRC and ACVD from other diseases.

We then focused on the VF genes encoding secretion systems and toxins and their pathogenic potential in ACVD and CRC. From the toxin’s perspective, 12 VF genes encoding colibactin in *Klebsiella pneumoniae* and two genes encoding heat-stable enterotoxin 1 and L-lysine 6-monooxygenase IucD in *Escherichia coli* were significantly enriched in patients with CRC, while only endotoxin genes participating in LPS and capsule biosynthesis were found in patients with ACVD.

We further analyzed the average abundance of VF genes in each type of secretion system separately (Figure S10). Remarkably, the type III secretion system VFs were enriched in many diseases, not limited to ACVD and CRC, whereas T6SS genes were more abundant in ACVD than in other diseases, implying their potential in inducing ACVD.

Given that the secretion systems in bacteria mediate bacterial-bacterial or host-bacterial competition by injecting diverse effectors, usually cytotoxic, into prokaryotic and eukaryotic cells^23^, we further analyzed the distribution of effectors in different groups (Figure S11). It was evident that different sets of effector genes were enriched in CRC and ACVD. As expected, many T3SS effectors were enriched in both CRC and ACVD patients. Importantly, we found the enrichment of one T6SS effector in the ACVD group, which supports our hypothesis that T6SS may play an essential role in the pathogenicity of ACVD.

In addition to fecal samples, we analyzed the respiratory tract metagenome of children, including 171 healthy children and 76 children with pneumonia. Overall, the diversity of VFs was significantly lower in healthy children’s respiratory tract microbiomes than in children with pneumonia (Figure S16). Specifically, adhesin-related genes in *Mycoplasma pneumoniae* were more abundant in children with pneumonia (Figure S17). There were significant differences in respiratory microbial virulomes between healthy children and children with pneumonia, probably due to the differences in oropharyngeal microbial diversity^24^.

### Gut virulome comparison in diabetes mellitus (DM) and gestational diabetes (GDM) with in-house sequenced datasets

We sequenced 150 fecal DNA samples from 50 healthy Chinese adults, 50 T2D (type 2 diabetes mellitus), and 50 T2D+CVD (cardiovascular disease) patients using Illumina sequencing technology. A total of ~ 11 Gb per sample was obtained. The sequencing statistics are summarized in Table S6.

We found that patients with type 2 diabetes and cardiovascular diseases (T2D+CVD) had a more diverse virulome than patients with type 2 diabetes (T2D) and healthy controls (Figures 4A and 4B). Nonmetric multidimensional scaling (NMDS) analysis showed a clear separation between patients with T2D and healthy controls (Figure 4D). Consistent with our observation that the VF abundances were higher than those in healthy controls (Figure 4C), we found that many VFs were significantly enriched in T2D+CVD and T2D samples compared with their healthy controls (Figure 4E). The LDA scores indicated that the abundances of autotransporter-related VFs were much more enriched in T2D, while adherence and T6SS were much more enriched in T2D+CVD. The most enriched VFs in T2D and T2D+CVD were derived from *Escherichia coli* and *Klebsiella pneumoniae.* Furthermore, we compared the abundance between mobile VFs and nonmobile VFs and found that nonmobile VFs were significantly higher than mobile VFs for each group (Figure S18).

**Figure 4.**
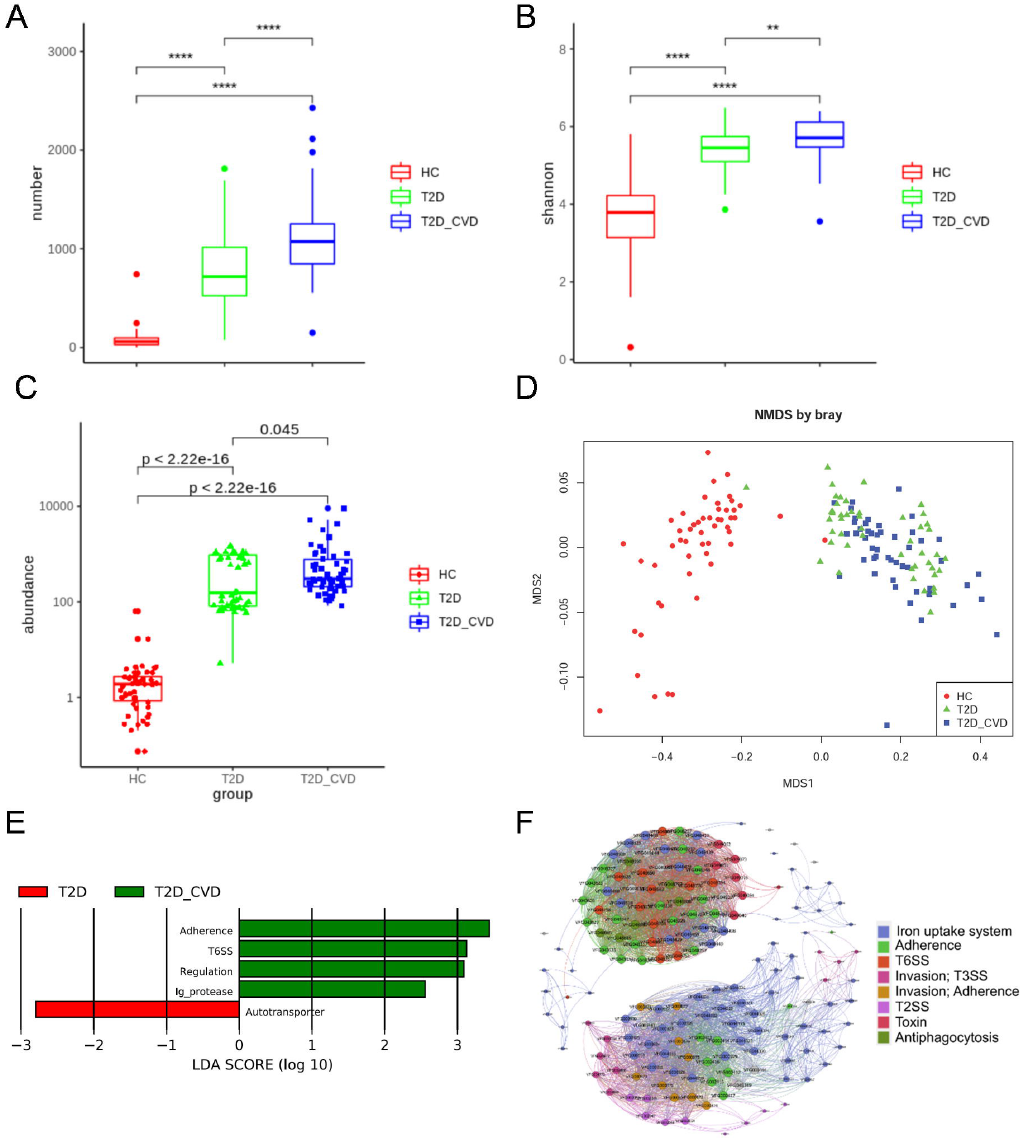
Patients with type 2 diabetes with cardiovascular diseases (T2D+CVD) had a more diverse virulome. (A) Boxplot of the number of VF genes present in each sample. (B) Boxplot of the Shannon diversity indexes of all samples based on the virulome. *p < 0.05, **p < 0.01, ***p < 0.001, ***p < 0.0001, Wilcoxon rank-sum test. (C) Comparison of the mean VF abundance. For each boxplot, the centerline represents the median, and the boxes correspond to the 25th and 75th percentiles; all data points are shown. (D) NMDS of Bray-Curtis dissimilarities showing the virulome. Bray-Curtis dissimilarities were calculated from the relative VF abundance profiles. The x-axis shows the first principal coordinate, and the y-axis shows the second principal coordinate. (E) Histogram of the LDA scores (log10) computed for VFs with differential abundance in the healthy, T2D, and T2D+CVD subjects. The LDA scores indicated that the abundances of autotransporter-related VFs were much more enriched in T2D, while adherence and T6SS were much more enriched in T2D+CVD. Most of the enriched VFs in T2D and T2D+CVD were derived from *Escherichia coli* and *Klebsiella pneumoniae.* (F) Network analysis demonstrating the co-occurrence patterns between VFs. The nodes are colored according to the VF genes, with each node representing a VF subtype. The size of each node is proportional to its number of connections. An edge is a strong (q > 0.6) and significant (P-value < 0.01) connection between nodes.

To indicate the relationship between VFs, we performed Spearman’s correlation analysis between VFs. The strong (q > 0.6) and significant (adjusted P value < 0.05) correlations between VFs are shown in Figure 4F. Two major modules were identified within the network. One module contained VFs relating to T6SS, toxin, antiphagocytosis, adherence, and the iron uptake system. The other module contained VFs relating to T3SS, T2SS, adherence, and the iron uptake system. The VF modules are of particular interest because they represent the functional relationship between VFs. They may provide a systems perspective at the community level.

In contrast, we did not find a more diverse virulome in patients with GDM than in their healthy controls (Figure S19). DM showed a significantly diverse virulome over their healthy controls, while GDM had no statistically significant diverse virulome. Therefore, GDM may represent transient DM, and the virulome appears to be relevant to DM pathogenesis but not GDM, although its underlying mechanisms are unknown.

### Selected samples of DM from short-read results confirmed by PacBio long-read sequencing

To experimentally confirm the presence of VF genes in the human gut microbiome, we sequenced 9 fecal DNA samples from 3 healthy Chinese adults, 3 patients with T2D, and 3 patients with T2D+CVD using PacBio single-molecule real-time (SMRT) long-read sequencing technology. A total of ~ 20 Gb per sample with an average subread length of 8 kb was obtained with the PacBio Sequel II system. The sequencing statistics are summarized in Table S7. The assembly of PacBio reads yielded 37 large CCs from 1 to 5 Mb in length, considered to be bacterial chromosomes. It also generated 149 CCs (73.4 to 947.4 kb) classified as plasmids and 5 CCs (54.4 to 12.2 kb in size) as phages.

Consistent with our findings using short-read sequencing, we found that many VF genes existed in fecal sample contigs from patients. The heatmap shows the VF distribution among the 9 human gut samples using SSI (Figure 5A). The mean numbers of VFs in T2D+CVD were significantly higher than those in the other two groups. Most of the VFs were derived from *Escherichia coli* and *Klebsiella pneumoniae*, consistent with Illumina sequencing observations. VF genes in the complete genome of the *Klebsiella pneumoniae* strain KP3037 are shown in Figure 5B. Specifically, two distinct gene clusters encoding T6SS were identified and confirmed by VRprofile^25^, a web-based tool for profiling virulence traits encoded within genome sequences of pathogenic bacteria. Mobile element-like genes, including genes involved in virulence and antibiotic resistance, were the major differences between strains.

**Figure 5.**
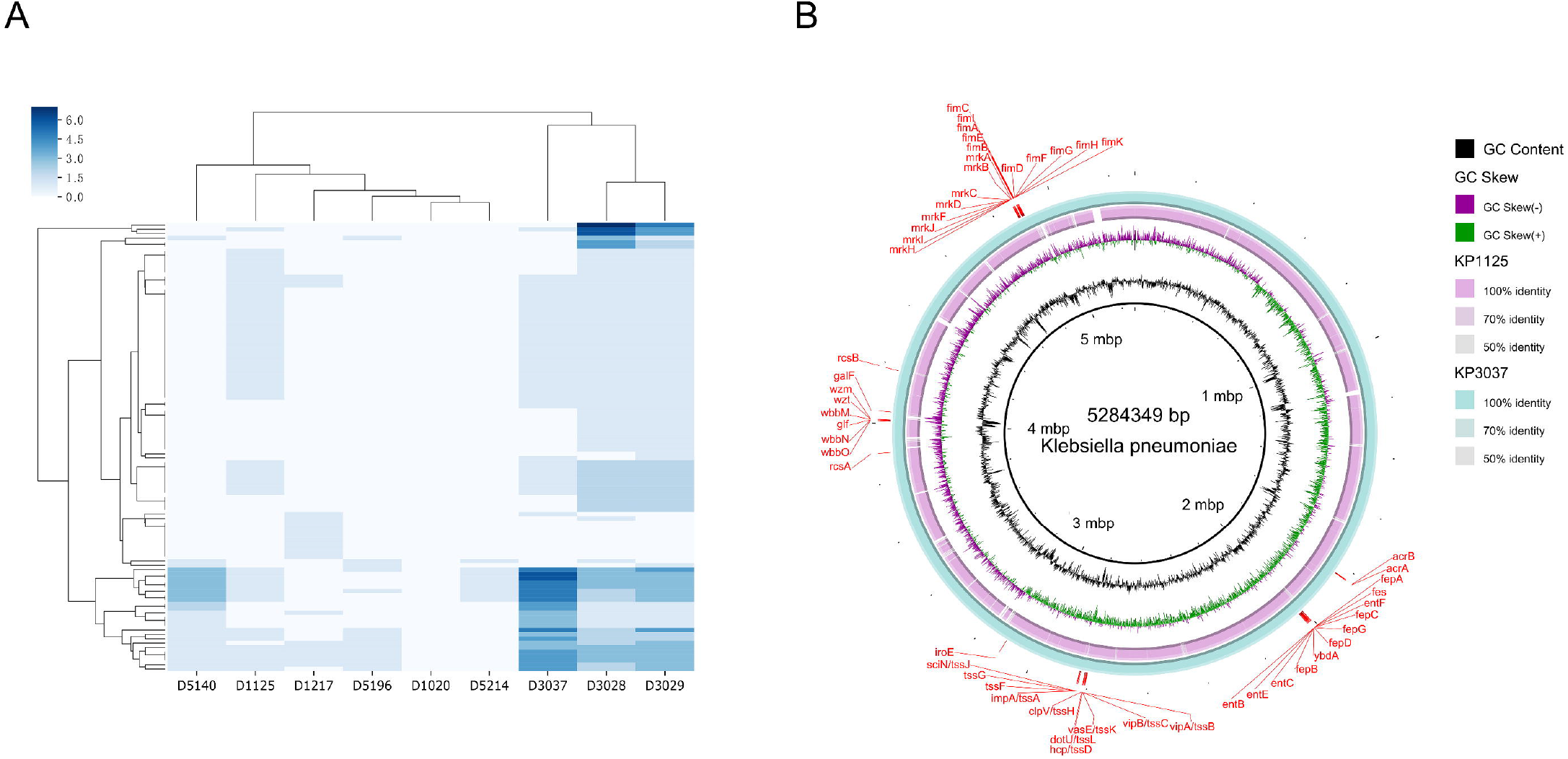
PacBio long-read sequencing confirmation of VF genes that exist in the contigs of fecal samples. (A) Heatmap shows the VF distribution among the 9 human gut samples using SSI. The mean numbers of VFs in T2D+CVD were significantly higher than those in the other two groups. Most of the VFs were derived from *Escherichia coli* and *Klebsiella pneumoniae*, consistent with Illumina sequencing observations. (B) BLAST ring image of the two complete genomes of *Klebsiella pneumoniae*. The *Klebsiella pneumoniae* strain KP3037 was used as the reference in the outermost ring. The two innermost rings represent the GC content of that area and the GC skew, respectively. The saturation of the color in these rings indicates identity by BLAST hit.

## Discussion

In this study, we conducted a comprehensive whole-body virulome analysis of the healthy human microbiota. We analyzed data from more than 4,000 samples across healthy and diseased subjects and 13 types of diseases from different metagenomic sequencing studies to characterize the disease-specific virulome. As the actual functions in the pathogenesis of predicted VF-related genes remain unclear, only experimentally verified VFs were involved in our study. We expanded the VF database termed VFGSSI and used species-specific sequence identity (SSI) inferred from the mean ANI values per species to precisely assign reads to virulence factors.

Our findings have substantially expanded our insight into the abundance and diversity of VFs in different body sites. Differences in the environmental conditions between different body habitats may be reflected in the microbiome and, consequently, the virulome. We observed a unique body-site virulome composition in this study. These findings illustrate that the healthy human microbiota, in general, beyond the gut microbiota, is a reservoir for virulence factors. This reservoir may serve as a mobile gene pool that facilitates VF transmission. The differences in eating habits, personal care, and lifestyles between men and women may lead to sex-specific differences in the composition of VF genes. Our results highlight the importance of sex-specific analysis when studying the human microbiome and virulome. New epidemiological studies are needed to evaluate the prevalence of potentially pathogenic bacteria carrying VFs in the healthy human body.

We hypothesized that the different diseases correspond to a specific virulome, especially in ACVD and CRC. Initially, the enrichment of genes encoding the type VI secretion system (T6SS) in *Klebsiella pneumoniae* was characteristic of the ACVD virulome, which was also discovered and then confirmed by PacBio’s single-molecule real-time (SMRT) sequencing in an independent dataset of the Diabetic Cardiovascular Complications cohort. T6SS is widely found in gram-negative bacteria, including *Bacteroidetes* and *Proteobacteria*, and is dedicated to mediating interbacterial antagonism and niche occupancy^26^. Recently, Verster *et al*. revealed the role of *Bacteroides fragilis* T6SS in mediating the gut microbe community^27^. Therefore, we assumed that the existence of T6SS genes might result in the overgrowth of *Klebsiella pneumoniae* in patients with CVD, which can explain why *Klebsiella pneumoniae* is enriched in CVD cohorts^28,29^. In addition, endotoxin (LPS) components of *Klebsiella pneumoniae* are another signature of ACVD. As it has been reported that low-grade chronic inflammation promotes the development of CVD^30^, the enrichment of LPS may lead to increased inflammation; therefore, it contributes to the development of ACVD.

In contrast to ACVD, patients with CRC exhibited an enrichment of genes encoding the secreted toxin colibactin (*clb*), which has been reported to be enriched in adenomatous polyposis (FAP) ^31^ and leads to CRC by inducing oncogenic mutations of enterocytes^32^. Although previous research has focused on the ability of colibactin production in *E. coli*, in our virulome analysis, *clb* genes were annotated to the genome of *Klebsiella pneumoniae.* Since colibactin genes are not present in intestinal pathogenic *E. coli* strains but are present in *E. coli* strains isolated from human feces^33^, it is reasonable that *clb* genes in *E. coli* were not found. In addition, the structure of *clb* is highly conserved among *Enterobacteriaceae*, including *Klebsiella pneumoniae*^34^. Thus, another assumption is that the carcinogenic potential is not limited to *E. coli* but may expand to other gut bacteria with *clb* gene clusters. Due to regional, temporal, and spatial differences, it is crucial to have matched healthy controls when studying the microbiome and virulome. Together, our results suggest that VF profiles are unique to each disease and that our approach for classifying virulomes can be applied more broadly.

Understanding the impact of virulence may provide new treatment options for microbe-related diseases. The differences in VF profiles across different body sites and disease types have significant implications for verifying the virulome and finding new antibacterial treatments. This work also provides a useful reference for future virulome studies in the human microbiome.

## Methods

### Dataset collection

A total of 1,497 metagenome datasets from habitats within the human skin, oral cavity, gut, and vagina from the HMP cohort^35^ were downloaded from the National Center for Biotechnology Information (NCBI) Sequence Read Achieve (SRA, http://www.ncbi.nlm.nih.gov/sra). Detailed information, including the sample ID, sequencing platform, read length, read number, data size, and accession numbers for each dataset, is shown in Supporting Information Table S8. The SRA datasets were converted to fastq using the fastq-dump module in the NCBI SRA Toolkit. We collected 2,712 samples from 13 types of diseases, including colorectal carcinoma (CRC)^36–39^, atherosclerotic cardiovascular disease (ACVD)^40^, inflammatory bowel disease (IBD)^3,41^, obesity^42^, hypertension^43^, Parkinson’s disease (PD)^44^, non-small cell lung cancer (NSCLC)^45^, hepatocellular carcinoma (HCC)^46^, gastric cancer (GC)^47^, cirrhosis^48^, melanoma^49,50^, renal cell carcinoma (RCC)^45^ and children with *Mycoplasma pneumoniae* pneumonia (MPP) ^24,51^. In total, we analyzed more than 4,000 metagenomic samples.

### DNA extraction and whole-genome sequencing

The total genomic DNA in fecal samples was extracted using a QIAamp PowerFecal DNA Kit, following the user manual. Total DNA was eluted in 200 μL of sterile water and stored at −20°C until use. A NanoDrop was used to measure the concentration and purity of the DNAs. Library preparation was carried out following the recommended protocol from BioScientific’s kit. Briefly, approximately 2 μg of DNA from each sample was used for fragmentation by Biorupter (high power: (15 s, on/90 s, off), six cycles) and end preparation by NEXT flex TM End-Repair. After PCR amplification (10 cycles), the library was purified using AMPure beads. Qubit was used to evaluate the quality and quantity of each library. For short-read sequencing of collected samples, whole-genome sequencing libraries were prepared using NexteraXT reagents (Illumina) and sequenced on an Illumina HiSeq X Ten platform. For long-read sequencing, SMRTbell libraries were sequenced on SMRT Cells (Pacific Biosciences) using magnetic bead loading and P4-C2 or P6-C4 chemistry.

### Virulence factor database curation

The VFDB (Virulence Factors of Bacterial Pathogens) database ^52^ is a comprehensive warehouse for deciphering bacterial pathogenesis. The VFDB (setA) core dataset comprises genes associated with experimentally verified virulence factors (VFs) for 53 bacterial species. PATRIC does not provide all the details for each VF and is not responsible for the original annotation. PHI-base focuses on plant pathogens. Although Victors includes VFs from bacteria, viruses, parasites, and fungi, VFDB focuses on human bacterial pathogens and contains more bacterial pathogens and experimentally verified VFs than Victors. This study downloaded the complete bacterial genomes from the NCBI server (accessed in Feb 2020), including 53 species of bacterial pathogens. Since the number of available genome sequences is unequal among different species, we randomly selected 100 genome sequences per species for ANI analysis and obtained averaged ANI values per species. For ANI calculations, the query organism’s genome is split into 1-kbp fragments, which are then searched against a reference organism’s whole genome. The average sequence identity of all matches having 60% overall sequence identity over 70% of their length is defined as the ANI between the two organisms^22^. To identify prophage-encoded VFs, we downloaded the complete virus genomes from the NCBI server (accessed in June 2020) and performed BLAST searches against the downloaded virus genome using the VFDB core dataset and the complete bacterial genomes (sequence identity 99%; coverage 99%).

We curated the gene annotation of experimentally verified VFs in the VFDB, which comprises 3,228 experimentally verified gene sequences from 53 species of bacterial pathogens. We identified VF gene sequences distributed across 74 species using a nucleotide identity cutoff value of 100% for the BLAST search against the chromosome sequences in the complete bacterial genomes. We performed intraspecies ANI analysis for each of the 74 species. The above-identified VF gene sequences with intraspecies ANI thresholds were used as the seeds to retrieve additional potential VF gene sequences from the complete bacterial genomes. Specifically, the complete bacterial genomes were subjected to local BLASTN against the VF gene sequences to hit potential VF sequences using species-specific sequence identity (SSI). The filtered hit sequences were extracted, and redundant sequences were removed from the whole database. A total of 56,913 VF gene sequences with SSI (VFGSSI) serve as a reference sequence for VF gene abundance calculation, of which 6,584 were mobile VFs and 50,329 were nonmobile VFs. The mobile VF gene sequences were identified using SSI thresholds for the BLAST search against the complete bacterial genome plasmid sequences.

### Metagenomic analysis

The virulome was determined first by aligning metagenomic reads to the dataset using BBMap with default parameters and then processed using a custom Python script to filter the mapped reads with the specific sequence identity inferred from the mean ANI values per species. For gene abundance calculation, the read counts aligned to this gene were normalized by the gene’s length and the total number of reads in the sample. We manually curated a pathogen list from a previous report^53^ to identify pathogenic and nonpathogenic strains.

MetaPhlAn2 ^54^ was used to perform taxonomic classification and profiling by mapping metagenomic reads against a library of clade-specific markers. PacBio sequencing reads were assembled by Canu ^55^. VirSorter ^56^ was used for the classification of CCs as phages. Categories 1, 2, 4, and 5 were considered phages, while categories 3 and 6 were excluded because they included false positives. PlasFlow ^57^ was used to identify plasmid-like contigs. Gene identification was performed on assembled sequences using MetaGeneMark^58^. The number of unique and shared VFs was calculated for the compared sample types, and Venn diagrams were drawn in Python using the Venn and matplotlib-venn packages.

### Statistical analysis

Principal coordinate analysis (PCoA) and nonmetric multidimensional scaling (NMDS) were performed to evaluate the differences in VF profiles among samples based on the Bray-Curtis distance of VF relative abundance. Permutational multivariate analysis of variance (PERMANOVA) between different groups was performed with adonis in vegan with a similarity index using 9999 permutations. LEfSe ^59^ analysis was used to identify discriminative VF types between groups. Diversity and heatmaps were prepared in R with vegan and ggplot2 packages.

## Author contributions

FL and BLZ conceived and designed the study; FL, WTD, YQG, XFS, YX, DMC,

XYF, YF, QX, NL, ZYL, JC, YNW collected and characterized the data; FL performed the data analysis; FL and WTD drafted the manuscript. All of the authors read and approved the final manuscript.

## Abbreviations

VFs: virulence factors
ACVD: atherosclerotic cardiovascular disease
IBD: inflammatory bowel disease
CRC: colorectal carcinoma
NSCLC: non-small cell lung cancer
HCC: hepatocellular carcinoma
GC: gastric cancer
PD: Parkinson’s disease
RCC: renal cell carcinoma
PCoA: principal coordinate analysis
NMDS: nonmetric multidimensional scaling.

## Acknowledgements

This work was supported in part by the Strategic Priority Research Program of The Chinese Academy of Sciences (XDB29020203) and the National Key R&D Program of China (2018YFC1603903 and 2018YFC1603803-2).

## Additional Files

### Additional file 1

**Figure S1. Schematic representation of the virulome analysis pipeline.** We curated the gene annotation of experimentally verified VFs in the VFDB, which comprises 3,228 experimentally verified gene sequences from 53 species of bacterial pathogens. We identified VF gene sequences distributed across 74 species using a nucleotide identity cutoff value of 100% for the BLAST search against the chromosome sequences in the complete bacterial genomes. We downloaded the complete bacterial genomes from the NCBI server (accessed on Feb 2020), including 74 species of bacterial pathogens. We performed intraspecies ANI analysis for each of the 74 species. The above-identified VF gene sequences with intraspecies ANI thresholds were used as the seeds to retrieve additional potential VF gene sequences from the complete bacterial genomes. Specifically, the complete bacterial genomes were subjected to local BLASTN against the VF gene sequences to hit potential VF sequences using species-specific sequence identity (SSI). The filtered hit sequences were extracted, and redundant sequences were removed from the whole database. The final VF gene sequences with SSI serve as a reference sequence for VF gene abundance calculation.

**Figure S2. Different body sites have distinct microbiomes.** (A) Boxplot of the Shannon diversity indexes of all samples from different body sites based on relative species abundance profiles. *p < 0.05, **p < 0.01, ***p < 0.001, ***p < 0.0001, Wilcoxon rank-sum test. (B) Principal coordinate analysis of Bray-Curtis dissimilarities showing the microbiome. Bray-Curtis dissimilarities were calculated from the relative species abundance profiles. The x-axis shows the first principal coordinate, and the y-axis shows the second principal coordinate.

**Figure S3. Comparison of mean VF abundance in the samples at six major body sites.** For each boxplot, the centerline represents the median, and the boxes correspond to the 25th and 75th percentiles; all data points are shown.

**Figure S4. Histogram of the LDA scores (log10) computed for VFs with differential abundance in different body sites.**

**Figure S5. Venn diagram showing the number of shared and unique VF genes among different body sites.** (A) Venn diagram of the four body sites. (B) Venn diagram of each pair of body sites. The number of shared and unique VF genes is shown. The shared and unique VF genes among the groups were investigated. We found that a total of 200 VF genes were shared among body sites. Interestingly, the oral cavity and skin shared more VFs (689 types) than those shared between the gut and oral cavity (443 types) or between the gut and skin (444 types).

**Figure S6. VF gene profiles were sex-specific and relatively stable over time.** Comparison of the total VF abundance between males and females in four major body habitats (A) and six major body sites (B). Comparison of the total VF abundance among samples from the same individuals over time in four major body habitats (C) and six major body sites (D). In the boxplots, the upper hinge represents the 75% quantile, the lower hinge represents the 25% quantile, and the centerline represents the median. Compared to men, women showed a higher VF abundance in the skin and gut (ANOVA, p < 0.05). Specifically, females had higher VF abundance in the anterior nares. The availability of longitudinal samples of different body sites over two years from individuals who did not take antimicrobial drugs afforded us the ability to investigate the stability of virulomes over time. There was no significant difference among samples from the same individuals except for the vagina, verifying that virulomes remained stable over a long period in different body habitats.

**Figure S7. Histogram of the LDA scores (log10) computed for VFs with differential abundance between males and females in the gut.**

**Figure S8. Histogram of the LDA scores (log10) computed for VFs with differential abundance between males and females in the oral cavity.**

**Figure S9. Histogram of the LDA scores (log10) computed for VFs with differential abundance between males and females in the skin.**

**Figure S10. Hierarchical clustering of the mean abundance of VFs encoding secretion systems for each type of disease.**

**Figure S11. Hierarchical clustering of the mean abundance of VFs encoding effectors of secretion systems for each type of disease.**

**Figure S12. Richness, Simpson, Shannon, and evenness diversity of VFs in ACVD samples.**

**Figure S13. Richness, Simpson, Shannon, and evenness diversity of VFs in CRC samples.**

**Figure S14. Richness, Simpson, Shannon, and evenness diversity of VFs in LC samples.**

**Figure S15. Histogram of the LDA scores (log10) computed for VFs with differential abundance in ACVD samples.**

**Figure S16. Richness, Simpson, Shannon, and evenness diversity of VFs in the children’s respiratory tract metagenome samples.**

**Figure S17. Histogram of the LDA scores (log10) computed for VFs with differential abundance in the children’s respiratory tract metagenome samples.**

**Figure S18. Comparison of mobile and intrinsic VF abundance. “Intrinsic VFs” are VFs located only on the bacterial chromosome. “Mobile VFs” are VFs** located on plasmids. Each dot represents a metagenome sample. For each boxplot, the centerline represents the median, and the boxes correspond to the 25th and 75th percentiles; all data points are shown.

**Figure S19. Richness, Simpson, Shannon, and evenness diversity of VFs in GDM samples.**

### Additional file 2

**Table S1. The number of VF gene sequences from each species in the dataset.**

**Table S2. Distribution of the number of sequences in the VF categories in the dataset.**

**Table S3. List of pathogens that can cause infections of the gastrointestinal tract and the diseases they cause.**

**Table S4. List of pathogens that cannot cause infections of the gastrointestinal tract and the diseases they cause.**

**Table S5. The top 10% (referring to the ratio of VF type numbers) of the most abundant VF types in each type of disease, which were considered the representative VFs, are summarized.**

**Table S6. The Illumina short-read sequencing statistics.**

**Table S7. The PacBio long-read sequencing statistics.**

**Table S8. Detailed information on 1,497 metagenome datasets from habitats within the human skin, oral cavity, gut, and vagina from the HMP cohort is summarized.**

